# Inverse occlusion, a binocularly motivated treatment for amblyopia

**DOI:** 10.1101/418871

**Authors:** Jiawei Zhou, Yidong Wu, Yiya Chen, Xiaoxin Chen, Yunjie Liang, Yu Mao, Zhimo Yao, Zhifen He, Fan Lu, Jia Qu, Robert F. Hess

**Affiliations:** School of Ophthalmology and Optometry and Eye hospital, Wenzhou Medical University, Wenzhou, Zhejiang 325003, PR China; McGill Vision Research, Department of Ophthalmology, McGill University, Montreal, Quebec, Canada H3G 1A4.

## Abstract

Recent laboratory finding suggest that short-term patching the amblyopic eye (i.e., inverse occlusion) results in a larger and more sustained improvement in the binocular balance compared with normal controls. In this study, we investigate the cumulative effects of the short-term inverse occlusion in adults and old children with amblyopia. A prospective cohort study of 18 amblyopes (10-35 years old; 3 with strabismus) who have been subjected to 2 hours/day of inverse occlusion for 2 months. Patients who required refractive correction or whose refractive correction needed updating were given a 2-month period of refractive adaptation. The primary outcome measure was the binocular balance which was measured using a phase combination task, the secondary outcome measures were the best corrected visual acuity which was measured with a Tumbling E acuity chart and convert to logMAR units and the stereo acuity which was measured with the Random-dot preschool stereotest. The averaged binocular gain was 0.11 in terms of the effective contrast ratio (z = −2.344, *p* = 0.019, 2-tailed Related samples Wilcoxon Signed Ranks Test). The average acuity gain was 0.14 logMAR equivalent (t(17) = 0.13, *p* < 0.001, 2-tailed paired samples t-test). The averaged stereo acuity gain was 253 arc seconds (z = −2.689, *p* = 0.007). Based on more recent research concerning adult ocular dominance plasticity, contrary to current practice, patching the amblyopic eye makes more sense; comparable acuity benefits, better compliance, better binocular outcome and applicable to adults as well as old children.

## 1. Introduction

Occlusion of the fixing eye has been the gold standard treatment for amblyopia ever since it was first introduced in 1743 by Conte de Buffon[1]. It has evolved over the years; partial rather than fulltime occlusion is now preferred and filters (i.e. Bangerter filters)[2], lenses (i.e. defocus, or frosted) and eye drops (i.e. atropine)[3, 4] have been used instead of opaque patches. It is effective in over 53% of cases in improving acuity in the amblyopic eye by more than 2 lines of logMAR acuity[5]. It does however leave something to be desired in a number of aspects. Compliance can be low[6] because it restricts school age children to the low vision of their amblyopic eyes for part of the day and also because of its psychosocial side-effects[7]. There is a relatively poor binocular outcome even though the acuity of the amblyopic eye is improved[8]. Its effects are age-dependent; effectiveness is much reduced for children over the age of 10 years old[9, 10]. Finally, it is associated with a 25% regression rate once the patch has been removed[11, 12]. It is effective but far from ideal. Interestingly, the basis of this widely accepted therapy is poorly understood. An explanation is often advanced in terms of “forcing the amblyopic to work” by occluding the fixing eye, which prompts the question, *what is stopping the amblyopic eye from working under normal binocular viewing?* This suggests that the problem of improving vision in the amblyopic eye, far from being simply a monocular issue, must have an underlying binocular basis (i.e., involving the fixing eye). Occlusion of the fixing eye must be, in some way, disrupting what is normally preventing the amblyopic eye from working when both eyes are open. Within the clinical literature this is known as suppression and one supposes that occlusion affects suppression in a way that is beneficial to the acuity of the amblyopic eye.

Recent laboratory studies have shown that short-term occlusion (i.e., 2 hours) is associated with temporary changes in eye dominance in normal adults. There are two things that are particularly novel about this new finding; first, these changes occur in adults and secondly, the eye that is patched becomes stronger in its contribution to the binocular sum. In other words, the eye balance is shifted in favour of the previously patched eye. This was first shown by Lunghi et al (2011)[13] using a binocular rivalry measure to quantify eye dominance. Since then there has been a wealth of information on this form of eye dominance plasticity in normal adults using a wide variety of different approaches[13-25]. Zhou et al (2013)[25] were the first to show that adults with amblyopia also exhibited this form of plasticity and that it tended to be of larger magnitude and of a more sustained form. They made the novel suggestion that it could provide the basis of a new therapeutic avenue for amblyopes in re-establishing the correct balance between their two eyes. Such a suggestion rests on the assumption that serial episodes of short-term occlusion can lead to sustainable long-term improvements in eye balance. The hallmark of this form of plasticity is that, once the patch has been removed, the patched eye’s contribution to binocular vision is strengthened. Zhou et al (2013)[25] suggested that to redress the binocular imbalance that characterizes amblyopia, it is the amblyopic eye that would need to be occluded, opposite to what has been in common practise for hundreds of years to improve the acuity in the amblyopic eye. Such a therapy, in principle, would be primarily binocular in nature (addressing the binocular imbalance as a first step), it would be expected to have much less compliance problems since it is not affecting the day to day vision of the patient and since it has been demonstrated in adults, it could be administered at any age. While this is well and good from a purely binocular perspective, the obvious question is how would occlusion of the amblyopic eye on a long-term basis (e.g., 2 hours or more a day for months) affect the acuity of the patched eye? The ethical basis for such interventions is not in doubt, as there is evidence indicating that such treatment is likely to be benefit rather than harm the vision of the amblyopic eye (including children). In the 1960s, so-called inverse occlusion was sometimes used in an attempt to treat eccentric fixation, which accompanies amblyopia in its more severe form. A review of these studies[26-30] leads to two conclusions; first, inverse occlusion did not make the amblyopia worse and second, acuity improved in the amblyopic eye in a percentage of cases. The percentage of patients whose vision improved was significantly less than that of classical occlusion in most[26, 29, 30], but not all[27, 28] studies, which could arguably be a consequence of the fact that studies on inverse occlusion were restricted to the more severe and resistant forms of amblyopia. Therefore, on the basis of recent laboratory studies on ocular dominance plasticity resulting from short term monocular occlusion[13-25] and previous clinical studies, on inverse occlusion designed to treat eccentric fixation[26-30], we have two expectations; first that inverse occlusion (i.e., occlusion of the amblyopic eye) should improve the binocular balance in patients with amblyopia and second, that improved acuity of the amblyopic eye should also be expected. Two additional benefits of this approach would be the expectation of better compliance, as the fellow eye is not occluded and its applicability to older children and adults, since ocular dominance plasticity occurs in adults.

To determine whether this radical departure from what is in common practice has any benefit, we studied the effects of inverse occlusion of 2 hours /day for 2 months on a group of 18 anisometropic and strabismic amblyopic teens and adults (10-35 years old), an age range where classical occlusion therapy has low compliance[31]. Our primary outcome measure was the binocular balance or ocular dominance. The second outcome measures were visual acuity and stereo acuity. The results suggest that this approach results in modest gains in both binocular balance and visual acuity within this older age group, no adverse effects were encountered.

## 2. Materials and Methods

### 2.1 Participants

Eighteen amblyopes with (n = 3) or without (n = 15) strabismus participated in our experiment. All of the patients were detected at 10 years or older or had failed with classical occlusion therapy (i.e., patching the fellow eye). Clinical details of patients are provided in Table 1. Observers wore their prescribed optical correction, if needed, in the data collection. Written informed consent was obtained from all patients, or from the parents or legal guardian of participants aged less than 18 years old, after explanation of the nature and possible consequences of the study. This study followed the tenets of the Declaration of Helsinki and was approved by the Ethics Committee of Wenzhou Medical University.

**Table 1.**
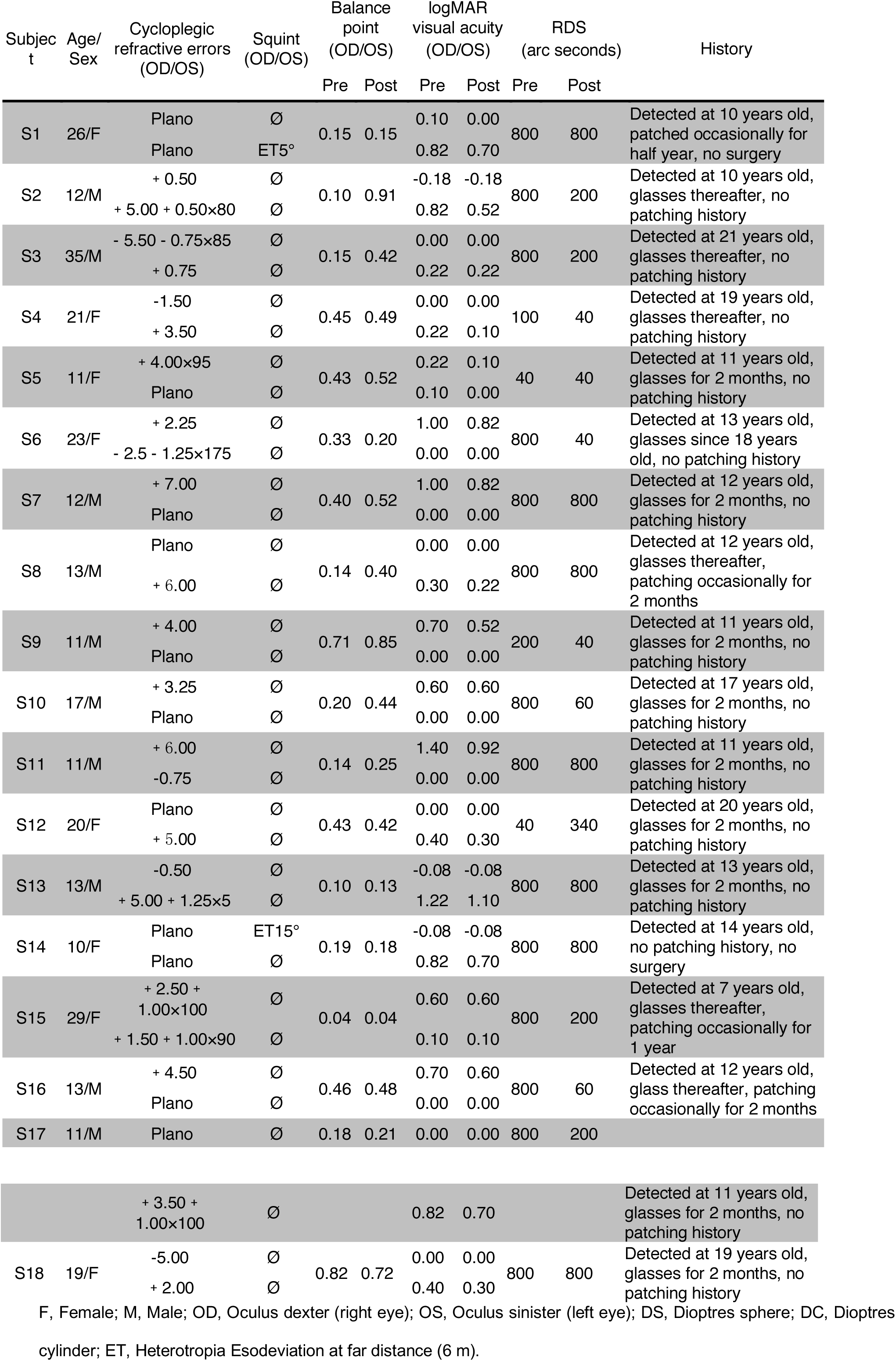
Clinical details of the participants.

### 2.2 Apparatus

The measures of binocular balance were conducted on a PC computer running Matlab (MathWorks, Inc., Natick, MA) with PsychToolBox 3.0.9 extensions[32, 33]. The stimuli were presented on a gamma-corrected LG D2342PY 3D LED screen (LG Life Science, Korea) with a 1920 × 1080 resolution and a 60 Hz refresh rate. Subjects viewed the display dichoptically with polarized glasses in a dark room at a viewing distance of 136 cm. The background luminance was 46.2 cd/m^2^ on the screen and 18.8 cd/m^2^ through the polarized glasses. A chin-forehead rest was used to minimize head movements during the experiment.

The measure of best-corrected visual acuity was using a Tumbling E acuity chart, the Chinese national standard logarithmic vision chart (Wenzhou Xingkang, Wenzhou, China), at 5 meters. This consists of E letters in 4 orientations on each line in a logarithmic progression from 20/200 to 20/10. The measure of stereo acuity was using the Random-dot preschool stereograms (RDS test; Baoshijia, Zhengzhou, China) at 40 cm. Strabismus angle was measured using the prism cover test.

### 2.3 Design

Patients’ binocular balance (balance point in the binocular phase combination task), visual acuity and stereo acuity were measured before and after two months of occlusion of the amblyopic eye for 2 hours/day (i.e., the inverse occlusion). For patients who required refractive correction or whose refractive correction needed updating (n = 9), a 2-month period of refractive adaptation was provided prior to the inverse occlusion study (Figure 1).

**Figure 1.**
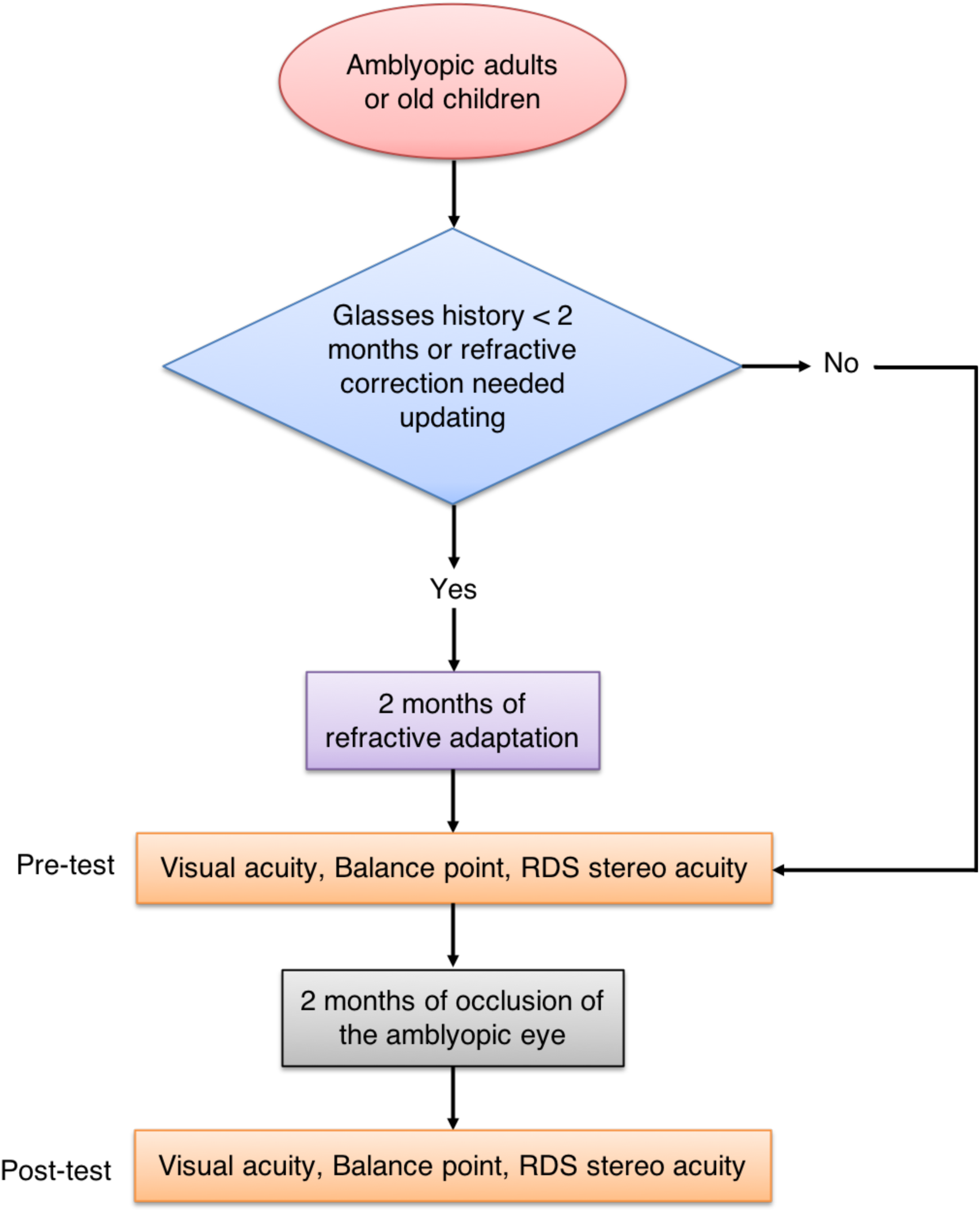
Experimental design. Eighteen amblyopes with (n = 3) or without (n = 15) strabismus participated in our experiment. Patients’ binocular balance (balance point in the binocular phase combination task), visual acuity and stereo acuity were measured before and after two months of occlusion of the amblyopic eye for 2 hours/day (i.e., the inverse occlusion). For patients who required refractive correction or whose refractive correction needed updating (n = 9), a 2-month period of refractive adaptation was provided prior to the inverse occlusion study.

Since this approach is different from that currently used (i.e., classical occlusion therapy), we were careful to conduct follow-up evaluations in accordance with the regulations from the Amblyopia Preferred Practice Pattern^®^ guideline (“PPP” 2017), P124: “If the visual acuity in the amblyopic eye is improved and the fellow eye is stable, the same treatment regimen should be continued”. In particular, we conducted weekly visits in the pilot study (in S1 to S13), rather than the 2 to 3 months that “PPP” recommends (P124 in “PPP”: “In general, a follow-up examination should be arranged 2 to 3 months after initiation of treatment “) to ensure that the acuity in the amblyopic eye did not deteriorate as a result of patching (Figure 2).

**Figure 2.**
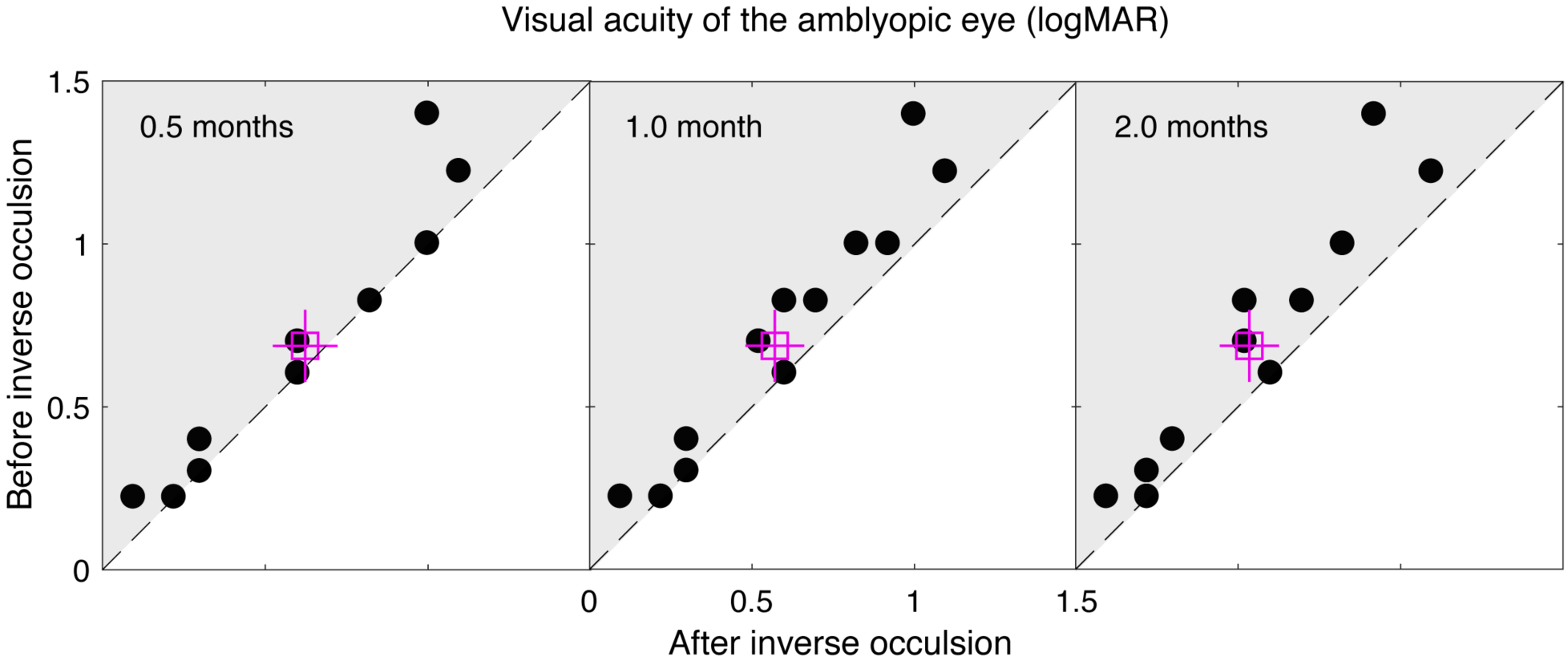
The change of amblyopic eye’s visual acuity after inverse occlusion. S1 to S13 participated in this pilot study. In each panel, each dot represents one patient. The open square represents the averaged results. Error bars represent standard errors. Data falling in the shaded area represents improvements; data falling on the sloping line represent no effect. Amblyopic eye’s visual acuity improved in 5 of the 13 patients after 2 weeks of treatment; in 9 of the 13 patients after 1 month of treatment; and in 11 of the 13 patients after 2 months of treatment. Fellow eye’s visual acuity was stable in all patients. No case of a deterioration of acuity in the amblyopic eye was recorded. The amblyopic eye’s visual acuity was significantly different at different follow-up sessions: F(3, 36) = 8.54, *p* < 0.001, 2-tailed within subject repeated-measure ANOVA.

We quantitatively accessed the binocular balance using a binocular phase combination paradigm[34, 35], which measures the contributions that each eye makes to binocular vision. The design was similar as the one we used in previous studies[36, 37], in which observers were asked to dichoptically view two horizontal sine-wave gratings having equal and opposite phase-shifts of 22.5° (relative to the center of the screen) through polarized glasses; the perceived phase of the grating in the cyclopean percept was measured as a function of the interocular contrast ratio. By this method, we were able to find a specific interocular contrast ratio where the perceived phase of the cyclopean grating was 0 degrees, indicating equal weight to each eye’s image. This specific interocular contrast ratio reflects the “balance point” for binocular phase combination since the two eyes under these stimulus conditions contribute equally to binocular vision. For each interocular contrast ratio (δ = [0, 0.1, 0.2, 0.4, 0.8, 1.0]), two configurations were used in the measurement so that any starting potential positional bias will be cancelled out: in one configuration, the phase-shift was +22.5° in the nondominant eye and −22.5° in the dominant eye and in the other, the reverse. The perceived phase of the cyclopean grating at each interocular contrast ratio (δ) was quantified by half of the difference between the measured perceived phases in these two configurations. Different conditions (configurations and interocular contrast ratios) were randomized in different trials, thus adaptation or expectation of the perceived phase would not have affected our results. The perceived phase and its standard error were calculated based on eight measurement repetitions. Before the start of data collection, proper demonstrations of the task were provided by practice trials to ensure observers understood the task. During the test, observers were allowed to take short-term breaks whenever they felt tired.

### 2.4 Stimuli

In the binocular phase combination measure, the gratings in the two eyes were defined as:

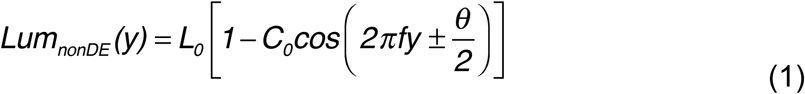

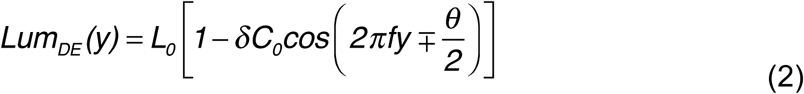

Where *L*_*0*_ is the background luminance; *C*_*0*_ is the base contrast in the nondominant eye; *f* is the spatial frequency of the gratings, *δ* is the interocular contrast ratio and *θ* is the interocular phase difference.

In our test, *L*_*0*_ = 46.2 cd/m^2^ (on the screen); *C*_*0*_ = 96%; *f* = 1 cycle/°; *δ* = [0, 0.1, 0.2, 0.4, 0.8, 1.0] and *θ* = 45°.

Surrounding the gratings, a high-contrast frame (width, 0.11°; length, 6°) with four white diagonal lines (width, 0.11°; length, 2.83°) was always presented during the test to help observers maintain fusion.

### 2.5 Procedure

We used the same phase adjustment procedure as used by Huang et al[35] for measuring the perceived phase of the binocularly combined grating. In each trial, observers were asked firstly to align the stimuli from the two eyes; they were then instructed to adjust the position of a reference line to indicate the perceived phase of the binocularly combined grating. Since the gratings had a period of 2 cycles corresponding to 180 pixels, the phase adjustment had a step size of 4 degrees of phase / pixel (2 cycles × 360 phase-degree / cycle / 180 pixels).

### 2.6 Statistical analysis

Data are presented as mean ± S.E.M unless otherwise indicated. Sample number (n) indicates the number of observers in each group, which are indicated in the figure. A one-Sample Kolmogorov-Smirnov Test was performed on each dataset to evaluate normality. A 2-tailed Related samples Wilcoxon Signed Ranks Test was used for comparison between nonnormally distributed datasets; A 2-tailed paired samples t-test was used for comparison between normally distributed datasets; A within subject repeated-measure ANOVA was used to evaluate the time effect of the inverse occlusion. Differences in means were considered statistically significant at *p* < 0.05. Analyses were performed using the SPSS 23.0 software.

## 3. Results

In the pilot study, we firstly conducted a 0.5-month of inverse occlusion (2 hours/day) in S1 to S13. We found that amblyopic eye’s visual acuity improvement in 5 of the 13 patients after 2 weeks of treatment, with no cases of acuity loss in the amblyopic eye. Visual acuity of the fellow eye was stable in all cases. We then extend the occlusion period to 1 month and 9 of 13 patients were found to exhibit small gains in visual acuity. No cases were recorded where the acuity of the amblyopic eye deteriorated. The Visual acuity of the fellow eye remained stable in all cases. We then extended the occlusion period to 2 months, and found that 11 of 13 patients showed small improvements in visual acuity in the amblyopic eye at that time. No patients exhibited a deterioration of function in the amblyopic eye and the visual acuity of the fellow eye remained stable (Figure 2). A within subject repeated-measure ANOVA verified that the amblyopic eye’s visual acuity was significantly different at these different follow-up sessions: F(3, 36) = 8.54, *p* < 0.001. This result clearly shows a dose-response relationship for the amblyopic eye in terms of visual acuity.

Since we could not have a control group who were denied any treatment, there is always the possibility that improvements in visual acuity measured at different time points are simply due to learning effects. To test this, we recorded the stability of acuity measured for the untreated fellow eye, as a similar learning effect should apply. In Figure 3, we plot the visual acuity gain as a function of treatment duration for the patched amblyopic eye and the unpatched fellow eye. There is an obvious difference between the two curves. A within-subject repeated-measure ANOVA, with eye and follow-up sessions as within-subject factors, verified that the visual acuity gain was significantly different between eyes (F(1,12) = 11.05, *p* = 0.006) and follow-up sessions (F(2,24) = 9.76, *p* = 0.001). The interaction between these 2 factors was also significant: F(2, 24) = 7.27, *p* = 0.003, indicating that the visual acuity gain of the amblyopic eye could not be accounted for by repeated testing alone.

**Figure 3.**
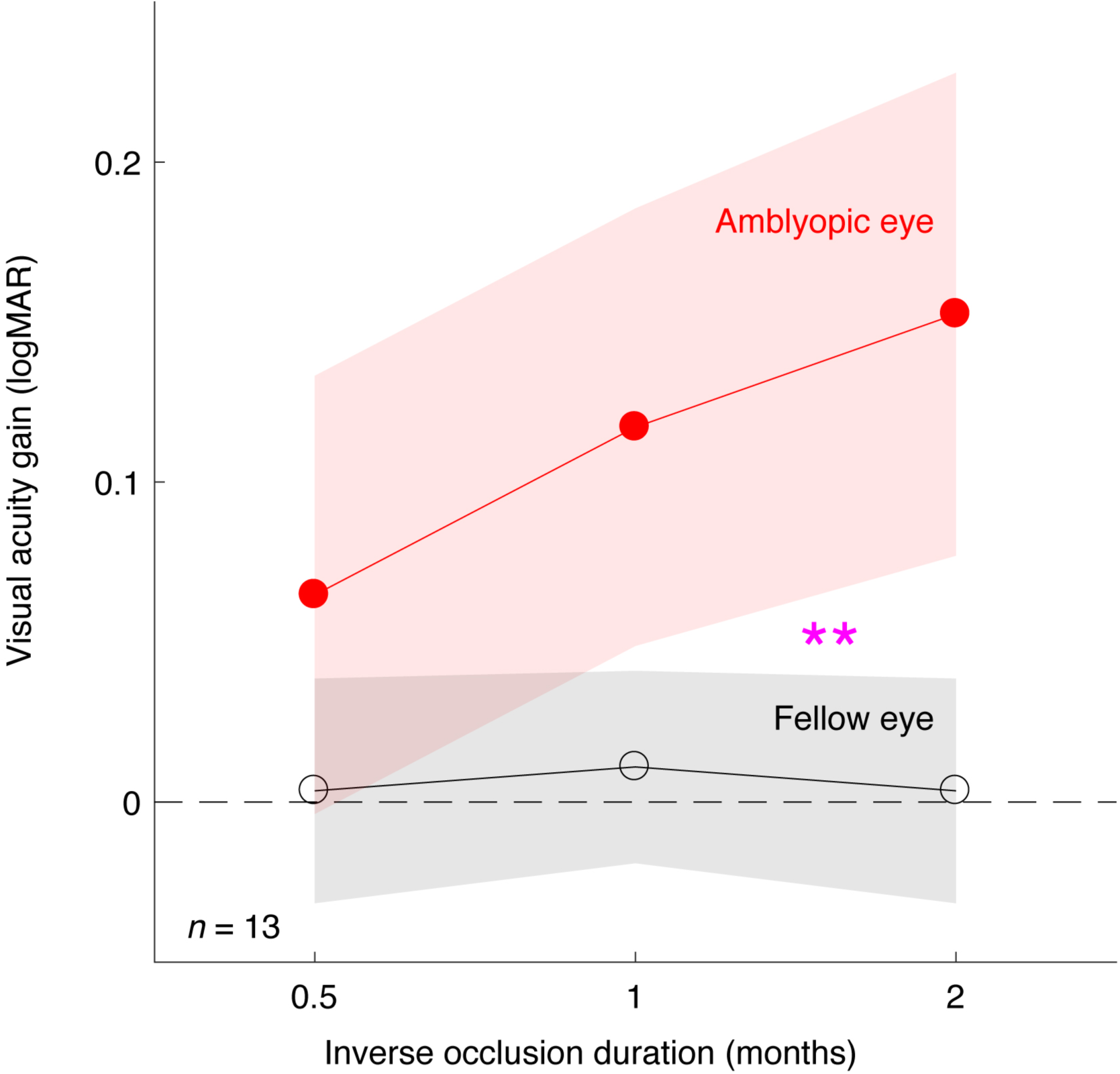
A dose-response relationship for the amblyopic eye. Averaged visual acuity gains of the amblyopic eye (filled circles) and the fellow eye (open circles) were plotted as a function of the inverse occlusion durations. The areas indicate the 95% confidence interval for mean. The two curves were significantly different (**): the interaction between eye and inverse occlusion duration was significant: F(2, 24) = 7.27, *p* = 0.003; 2-tailed repeated-measure ANOVA.

Once we had shown that inverse occlusion can be undertaken in a safe fashion, we added 5 additional patients (S14 to S18) to the original study cohort of 13 (S1 to S13). These additional patients followed the same protocol as the original thirteen (S1 to S13), but visual functions were only measured before and after 2 months of treatment. A summary of the main result for all the 18 patients is shown in Figure 4 for the measures of ocular balance, visual acuity and stereo acuity. Measurements before and after 2-month of treatment are plotted against one another. In term of ocular balance, the measure used is the interocular contrast that is required to achieve a binocular balance. By binocular balance we mean that the contributions of each eye’s input are equal at the site of binocular combination. For normals with equal eye balance, the effective contrast ratio would be unity. Values below unity indicate a shift in ocular dominance towards the fixing eye. Data falling on the sloping diagonal line represents no change from treatment whereas data falling in the shaded regions represents an improvement in binocular function (Figure 4A).

**Figure 4.**
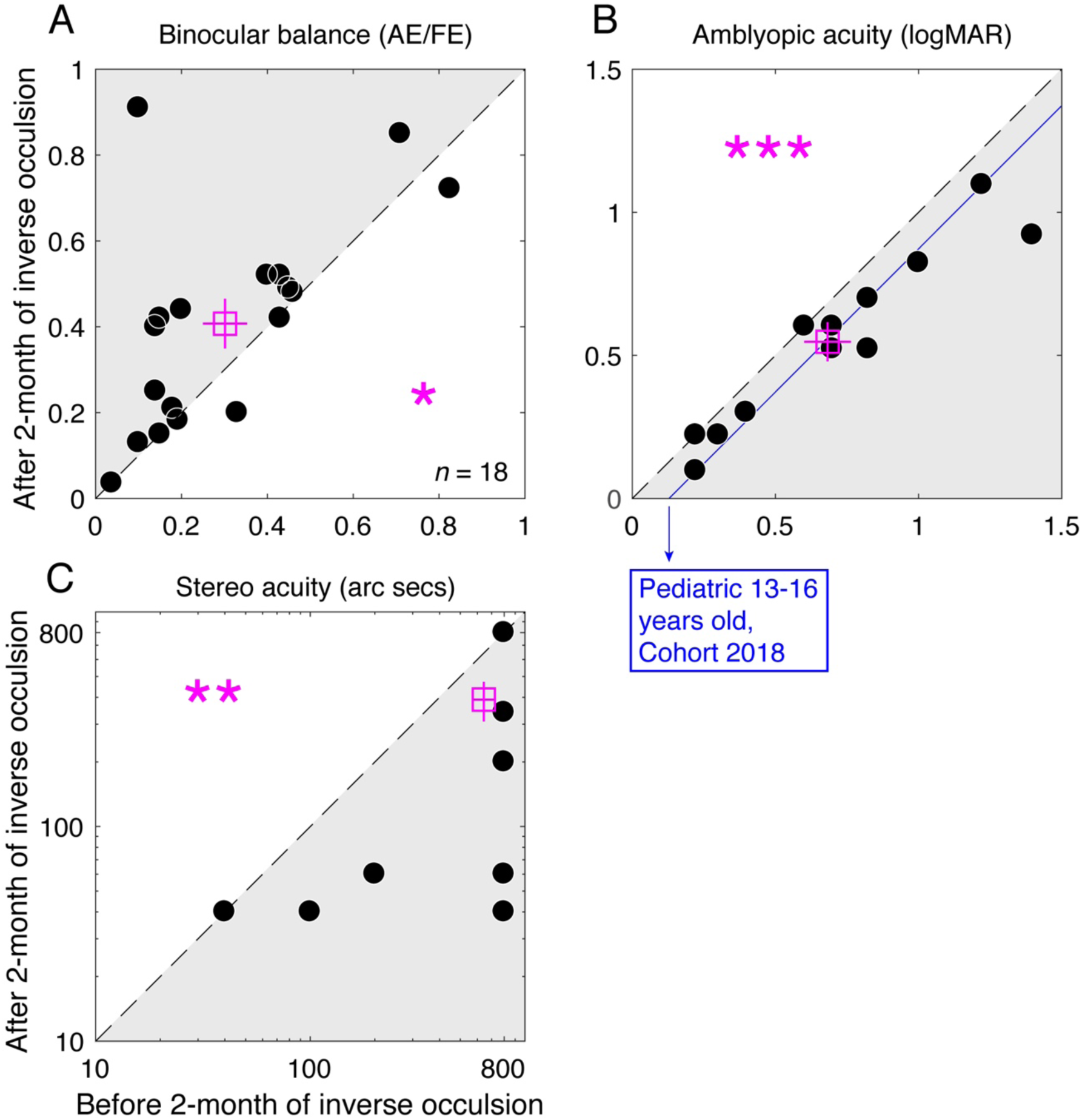
Visual outcomes after two months of occlusion of the amblyopic eye for 2 hours/day. Eighteen amblyopes (S1 to S18; 10 to 35 years old), with (n=3) or without (n=15) strabismus, participated. For patients who required refractive correction or whose refractive correction needed updating (n = 9), a 2-month period of refractive adaptation was provided before the inverse occlusion. A. Binocular balance was measured with the binocular phase combination task and expressed as the interocular contrast ratio (amblyopic eye / fellow eye) when the two eyes are balanced. The binocular balance increased from 0.30 ± 0.052 (Mean ± S.E.M.) to 0.41 ± 0.058. ‘*’: z = −2.344, *p* = 0.019, 2-tailed Related samples Wilcoxon Signed Ranks Test. Error bars represent standard errors. Data falling in the shaded area indicate patients whose two eyes were more balanced; data falling on the sloping line represent no change. B. Visual acuity was measured with a Tumbling E acuity chart in logMAR units. The visual acuity improved from 0.70 ± 0.085 (Mean ± S.E.M.) to 0.56 ± 0.070. ‘***’: t(17)=0.13, *p* < 0.001, 2-tailed paired samples t-test. Error bars represent standard errors. Data falling in the shaded area represents better visual acuity; data falling on the sloping line represent no change. The blue line indicates a 0.13 logMAR visual acuity improvement observed from a recent cohort study from the PEDIG group based on 2 hours daily of classical patching treatment for 16 weeks in children aged 13 to 16 years with amblyopia[38]. C. Stereo acuity was measured with the Random-dot stereograms. Stereo acuity of 800 arc secs was assigned for patients (14/18) whose stereo acuity was too high to be measured. The stereo acuity improved from 643.3 ± 71.48 (Mean ± S.E.M.) to 390 ± 81.48. ‘**’: z = −2.689, p = 0.007, 2-tailed Related samples Wilcoxon Signed Ranks Test. Error bars represent standard errors. Data falling in the shaded area represents better stereopsis; data falling on the sloping line represent no change.

Amblyopes exhibit a range of binocular balances ranging from less than 0.04 to 0.82 (Figure 4A). Inverse patching of 2 hours/day for 2 months improves some more than others. Six subjects showed no improvement, the other patients showed varying levels of improvement, meaning that their amblyopic eye was contributing more to binocular vision. Overall, the averaged improvement was a 0.11 change (0.30 ± 0.052 (Mean ± S.E.M.) to 0.41 ± 0.058) in the effective contrast ratio (Square symbol), which was significant based a 2-tailed Related samples Wilcoxon Signed Ranks Test: z = −2.344, *p* = 0.019. Our patients exhibited a range of acuity deficits ranging from less than 0.22 to close to 1.40 logMAR (Figure 4B). As expected, the acuity improvements were of varying degrees. Three patients showed no improvement at all, while all the other patients did exhibit improvements to varying degrees (shaded area). The averaged improvement (solid symbol) was 0.14 logMAR (from 0.70 ± 0.085 to 0.56 ± 0.070), which was significant based on a 2-tailed paired samples t-test: t(17)=0.13, *p* < 0.001. This magnitude of acuity gain is similar to the results of a recent PEDIG study using classical occlusion of the same duration (i.e. 2 hours/day for 16 weeks) in patients of a similar age range[38]. The averaged stereo acuity gain was 253 arc seconds (z = −2.689, *p* = 0.007, 2-tailed Related samples Wilcoxon Signed Ranks Test). This is a very conservative estimate because 14/18 patients had stereo acuities outside of our measurement range and were conservatively scored at 800 arc secs, the largest disparity tested. This means that the true stereo acuity gain could be larger than 253 arc seconds.

These changes in binocular balance, visual acuity and stereo acuity are modest but still impressive considering the fact that the period of occlusion was relatively short (2 hours), the duration of the treatment limited to 2 months and it involved an older age group. One interesting finding is that the improvements in balance and visual acuity are not significantly correlated (*p* = 0.76, Spearman’s correlation), so it is unlikely they have a common basis.

These improvements are long lasting as we have followed four patients (S12, S14, S16 and S17) for 1 month and one (S9) for 5.5 months after finishing the 2-month of reverse occlusion regime, which showed that the outcomes were sustained (Figure 5).

**Figure 5.**
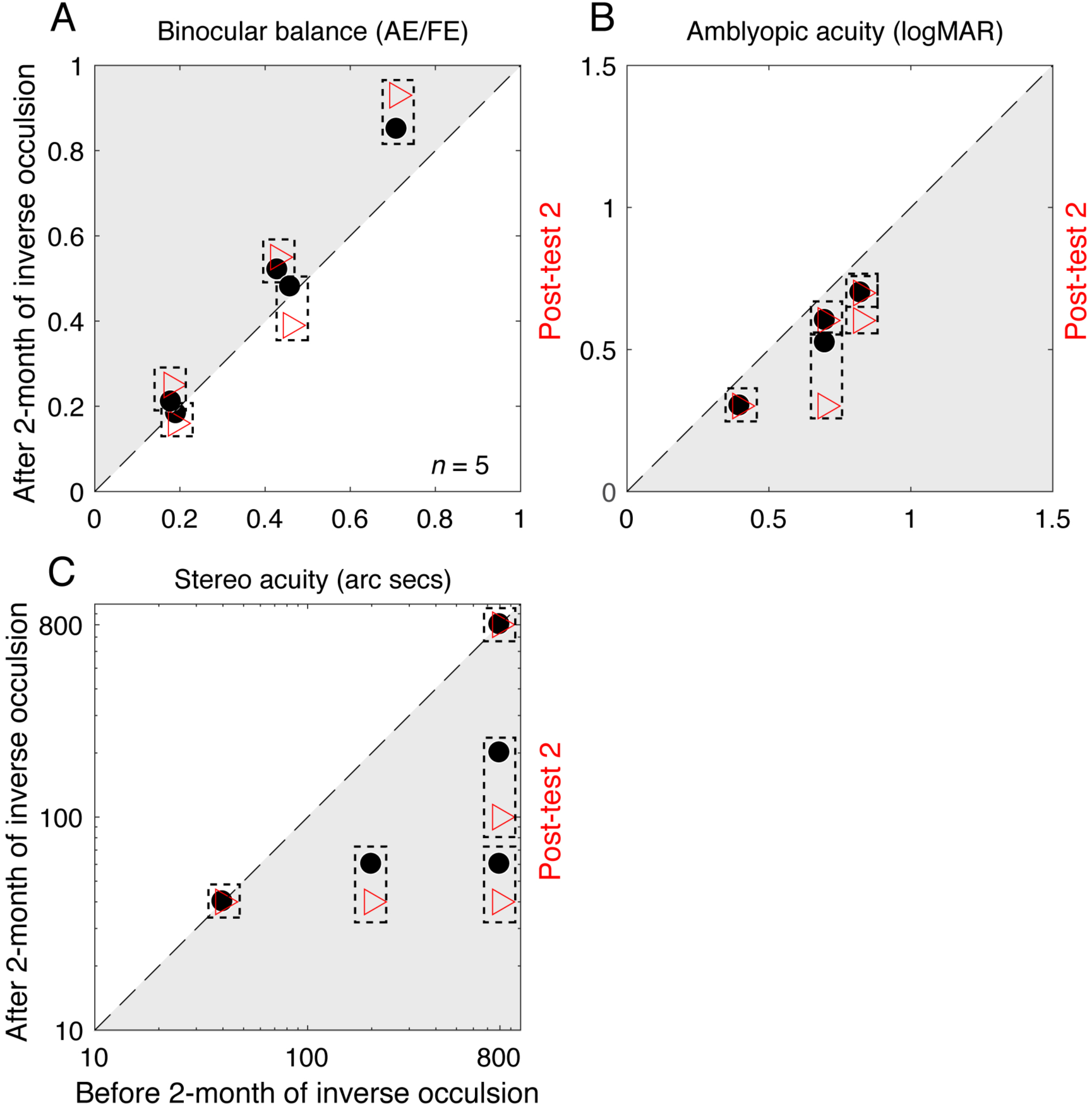
The visual outcomes could be sustained after finishing the 2-month of inverse occlusion. Four patients (S12, S14, S16 and S17) were re-measured at 1 month and one (S9) at 5.5 months after the completion of the 2-month of reverse occlusion regime. Their results that were measured immediately after the 2-month of inverse occlusion are marked as black dots; their results that were measured later are marked as red triangles. The corresponding results for each patient are marked using the dashed rectangle.

In our study, patients’ age ranged from 10 years old to 35 years old. Interestingly, all patients who were younger than 14 years old had visual acuity gain. However, for patients older than 14 years old, only 62.5% of them had a visual acuity gain. A Spearman correlation analysis showed that there was a positive correlation between the improvement in visual acuity of the amblyopic eye and the patients’ age, i.e., the younger the patients the more the visual acuity gain (Rho = 0.534, *p* = 0.022). The correlations between patients’ age and the binocular balance gain or the RDS stereo acuity gain were not significant (*p* > 0.3).

The refractive correction needed updating in half of the patients (n = 9), and a 2-month period of refractive adaptation was provided before inverse occlusion was commenced. Even though the acuity gains from optical treatments have been shown to be modest after 5-6 weeks of refractive adaptation[39], since those observations were in a much younger age group, there could still be an argument that our findings were due to the refractive correction *per se occurring after our 8-week period*, rather than the inverse occlusion. To assess this, we divided our patients into two subgroups, i.e., those who required refractive adaptation (n = 9) and those who did not (n = 9). We found no significantly different of visual outcomes in these two subgroups, in terms of the improvement of the amblyopic eye’s visual acuity (Z = −0.72, *p* = 0.49), binocular balance (Z = −0.13, *p* = 0.93) and stereo acuity (Z = −1.80, *p* = 0.09). Thus, there is no basis for believing that the gains were show here as the result of inverse occlusion where significantly impacted by refractive adaptation gains in visual acuity occurring beyond our 8-week refractive adaptation period.

## 4. Discussion

The rationale for this study comes from the recent findings on ocular dominance plasticity in normal and amblyopic adults[13-25], the finding that short term patching results in a strengthening of the contribution of the previously patched eye to binocular vision. This study, which applies this to amblyopia, raises three interesting issues that are relevant to the treatment of amblyopia. First, it highlights just how poor our understanding of the basis of classical occlusion therapy is. How is it that acuity improves in the amblyopia regardless of which eye is occluded? This does not just come from this study; there is a literature on the acuity improvements that occur as a result of inverse occlusion. While in most cases these improvements are much less than that of classical occlusion, there are studies[27, 28], where it is comparable to that of classical occlusion. The standard explanation of occluding the fixing eye to “forcing the amblyopic eye to work” is untenable. What is preventing the brain using information from the amblyopic eye under normal viewing conditions? Whatever it is, occlusion must be preventing (i.e., disinhibiting) it from operating. The problem must be essentially binocular in nature, which is why it is not critically dependent on which eye is occluded to disrupt the anomalous interaction. We would normally think about this anomalous binocular interaction as a suppression of the amblyopic eye by the fellow eye, but on the basis of the occlusion of either eye being effective, it may be better to think of suppression as simply a reflection of a binocular imbalance. Recent psychophysics [40] and animal neurophysiology[41] suggest that the problem is not because the inhibition from the fixing to the amblyopic eye is greater but because the matching inhibition from the amblyopic eye is less. It is due to a net imbalance in interocular inhibition. The resulting net imbalance can be disrupted by occluding either eye and it’s the duration of relief from this imbalanced binocular inhibition that may result in an acuity benefit for the amblyopic eye.

Ocular dominance plasticity in normals is an all-or-none, homeostatic process and would not be expected to have accumulated effects over time[42]. In amblyopes, ocular dominance plasticity has different dynamics, being much more sustained[25]. The present results suggest also that it can exhibit accumulated effects in amblyopes that result in long lasting changes in eye balance. These sustained changes are however modest in size and it will be necessary to explore how the magnitude of this effect can be increased for it to have significant binocular benefits. Future directions could involve RCT studies with large number of patients and longer durations of occlusion, potentially with pharmacological enhancement using dopaminergic[43], serotinergic[44] or cholinergic modulations[45] or the combination of binocular training procedures[46-50] and short periods of inverse occlusion.

The finding that the binocular balance and the monocular acuity improvements from inverse patching are not correlated suggests that a simple explanation in terms of reduced suppression is not viable. The two visual improvements are likely to have separate causes and possibly involving different sites in the pathway. The acuity improvement for the amblyopic eye is not dependent on which eye is occluded, as shown here (Figure 4B), but the direction of the binocular balance change is dependent on which eye is occluded[13, 25]. This distinction between binocular balance and monocular visual acuity is an important one and should be incorporated into future clinical treatment studies. Finally, apart from the additional benefit of a better binocular balance, its applicability to older children and adults should not be underestimated, nor should the better compliance that should follow from the patching of the amblyopic rather than the fixing eye. Application to younger children would necessitate weekly visits to ensure that the acuity in the amblyopic eye did not deteriorate as a result of patching.

### 4.1 Relevance of a recently published study

During the writing up of this paper, another study was posted on bioRxiv that is highly relevant and supportive of the present approach (Lunghi et al (2018); doi: https://doi.org/10.1101/360420). Lunghi et al (2018) undertook a comparable inverse occlusion study in adults based on the similar notion that patching of an eye can improve its contrast gain subsequently, a result that they originally showed in normal humans[13] and we originally demonstrated in humans with amblyopia[25]. However, Lunghi et al (2018) incorporated physical exercise as well as inverse occlusion and argue, based on a non-exercise control, that the combination of these two factors results in larger improvements when treating amblyopia. This in turn was based on their previous finding that exercise can enhance plasticity in normal adults ([18], but also see [23]). This published study and the current one both suggest that inverse occlusion can provide long term benefits in visual acuity, stereopsis and sensory balance. Lunghi et al find that six 2-hour sessions of inverse occlusion (*n* = 10) combined with exercise results in a visual acuity improvement of 0.15 ± 0.02 logMAR, whereas in our initial experiment of 13 patients (S1 to S13), we find a comparable improvement (0.15 ± 0.03 logMAR) after 2 months of 2hrs a day patching. The shortest treatment duration that we used involved 14 days of 2 hrs/day inverse occlusion and the acuity improvement was 0.06 ± 0.03 logMAR, similar to that found by Lunghi et al for their non-exercise control (0.06 ± 0.01 logMAR). The exercise enhanced protocol seems to be beneficial over the short treatment duration tested (i.e., 6 x 2 hrs periods). It will be interesting for future studies to compare the duration-response curves for inverse occlusion with and without exercise to know if they are parallel or whether they converse at longer treatment durations.

### 4.2 Shortcoming of the present study

These are pilot results, which we hope will help power larger RCTs on the potential benefits of inverse occlusion. The acuity results are modest and while they are comparable to those found for classical patching for the same short treatment duration[38], it would need to be shown that longer treatment durations result in at least the same extra benefits that has been shown for classical occlusion[51]. The binocular balance changes, while in the right direction are quite modest in magnitude and it would need to be shown that longer treatment durations would result in stronger accumulated effects. If this can be shown, inverse occlusion would carry an additional binocular benefit over that of classical occlusion. Finally, no adverse effects were found from this relatively short treatment duration in this older age group, future studies would need to assess this for longer treatment durations and younger age groups.

## 5. Conclusions

We conclude that patching the amblyopic eye is safe for adults as well as old children with amblyopia, and can result in recovery of visual acuity of the amblyopic eye and binocular visual functions.

## Data Availability

All data concerning this study is available within the manuscript. Detailed data is available upon request to the first author.

## Conflicts of Interest

The authors declare no competing interests.

## Funding Statement

This work was supported by the National Natural Science Foundation of China grant NSFC 81500754, the Qianjiang Talent Project (QJD1702021), the Wenzhou Medical University grant QTJ16005 and a grant from the Ministry of Human Resources and Social Security, China to JZ, and Canadian Institutes of Health Research Grants CCI-125686 and 228103, and an ERA-NET Neuron grant (JTC2015) to RFH. The sponsor or funding organization had no role in the design or conduct of this research.

